# An ensemble learning method for joint kernel association testing and principal component analysis on multiple kernels

**DOI:** 10.1101/2025.02.07.637007

**Authors:** Hyunwook Koh

## Abstract

In high-dimensional omics studies, researchers often conduct kernel association testing to powerfully detect the relationship of the genetic or microbial composition with human health or disease. Especially, in human microbiome studies, its dimension reduction analysis follows to visually represent complex microbiome data in a simple two- or three-dimensional coordinate space. However, various kernels exist, and they produce all different outcomes; hence, it is hard to interpret them all consistently. Then, omnibus testing has recently been a subject of intense investigation for a unified and powerful statistical inference. However, current omnibus tests are purely a test for significance producing only a *P*-value as their outcome with no related dimension reduction and visualization approach; hence, their utility is still limited. In this paper, I introduce an ensemble learning method, named as enKern, for joint kernel associating testing and principal component analysis on multiple kernels. enKern is based on a weight learning scheme that leverages complementary contributions from multiple kernels for powerful performance for various association patterns. I show that applying the weights to individual test statistics or individual kernels is equivalent, which in turn enables a visualization in a reduced dimensional coordinate space based on the weighted kernel to be matched with its original significance testing scheme. I demonstrate its use for human microbiome *β*-diversity analysis. I also demonstrate its outperformance in validity and power through simulation experiments. enKern is freely available in R computing environment at https://github.com/hk1785/enkern.

## 1 Introduction

In high-dimensional genome or microbiome studies, researchers often conduct kernel association testing (KAT) to powerfully detect the relationship of the genetic or microbial composition with human health or disease. The most foundational and widely used methods are sequence kernel association test (SKAT) [1, 2] in genomics, and microbiome regression-based kernel association test (MiRKAT) [3] in metagenomics. The advantages of KAT are first in an improved statistical power that is achieved by combining multiple association signals from multiple underlying variants while robustly capturing their correlation structures. Thus, KAT is especially useful for human genome or microbiome studies, where most genetic or microbial variants are rare variants in a complex correlation structure (e.g., genetic architecture, linkage disequilibrium, phylogenetic or taxonomic relationship) that tend to make weak individual effects, but a strong collaborative effect. Other advantages are in line with its regression-based modeling that facilitates (i) covariate adjustments that are needed to control for potential confounders (e.g., age, sex), and (ii) extensions to various types of response variables and/or study designs [4, 5, 6, 7, 8, 9, 10, 11].

In human genome studies, it has been widely accepted to improve statistical power at a cost of limited interpretation because it is too hard to discover causal variants. The reason is, for example, because of the requisite multiple testing correction for a couple of million variants given in the data that are, however, again only a small portion of the whole genome that contains billions of variants. Hence, it is likely that causal variants are not even contained in the data. However, in human microbiome studies, the situation is not that much harsh, and researchers intensely seek interpretable methods. For example, they often visualize complex microbiome composition for different study subjects in a simple two- or three-dimensional coordinate space using a dimension reduction method [12, 13, 14], such as principal component analysis (PCA) or multidimensional scaling [15, 16, 17]. However, in reality, various kernels exist, and they produce all different outcomes in significance and visualization; hence, it is not possible to interpret them all consistently. The situation is also that they reflect different views on the similarity or relatedness, and there is no kernel that is just right or wrong, or superior to the others in all contexts; hence, we cannot select a single best kernel to use in advance. In tradition, example kernels are linear, polynomial, gaussian kernels, and so forth; yet, SKAT [1, 2] employed the linear, quadratic and identity-by-state kernels, and their weighted versions, while MiRKAT [3] employed ecological kernels that are constructed by transforming ecological distance measures, also known as *β*-diversity indices, to similarity measures.

Recently, omnibus testing has been a subject of intense investigation for a unified and powerful statistical inference across different outcomes from multiple association tests. Especially, MiRKAT [3] employed so-called the minimum *P*-value approach for its omnibus test, named as OMiRKAT [3]. The minimum *P*-value approach is not a cherry-picking or fishing approach [18] that reports only the outcome that makes the minimum *P*-value that is the strongest evidence of significance. Of course, such a cherry-picking or fishing approach is not valid in significance testing with inflated type I error rates [18]. The minimum *P*-value approach takes the minimum *P*-value from multiple association tests as its test statistic [19], while building its null distribution through resampling. Here, the minimum *P*-value reflects the strongest signal among multiple association tests, while the resampling method captures complex correlation structures with no requisite parametric assumption. Hence, the minimum *P*-value approach maintains a high statistical power especially for a high sparsity level, where only few tests make strong signals, while robustly controlling type I error rates even when the underlying population distributions are highly skewed and correlated. Moreover, the minimum *P*-value approach can flexibly apply to kernel association tests [3, 8, 9, 10, 11], non-kernel association tests [20, 21, 22] or both kernel and non-kernel association tests [23, 24]. However, as with other omnibus tests [25, 26, 27, 28], its limitation is that it is purely a test for significance producing only a *P*-value as its outcome; hence, its interpretation is substantially affected. In methodological aspects, how to conduct kernel association testing and visualization jointly on multiple kernels has not been yet addressed.

In this paper, I introduce an ensemble learning method, named as enKern, for joint KAT and PCA on multiple kernels. enKern is based on a weight learning scheme that leverages complementary contributions from multiple kernels, instead of focusing only on the strongest signal as in the minimum *P*-value approach, for more robustly powerful performance for various association patterns. I show that applying the weights to individual test statistics or individual kernels is mathematically equivalent to formulate its weighted test statistic, which in turn enables dimension reduction and visualization based on the weighted kernel to be perfectly matched with its original significance testing scheme. Besides, a fast and flexible non-parametric permutation method is also feasible, which promises potential extensions to more kernels in a computationally efficient manner, and also can apply flexibly to various regression models.

While the application of enKern can range much broader, I demonstrate its use for human microbiome *β*-diversity analysis [12, 29, 3] through the reanalysis of two public real microbiome datasets: (1) to see the disparity in upper-respiratory-tract microbiome by cigarette smoking [30]; and (2) to see the disparity in oral microbiome by gingival inflammation and cytokine levels [31]. I also compare enKern with OMiRKAT [3] through simulation experiments, and show that (1) both enKern and OMiRKAT correctly control type 1 error rates, but (2) enKern is more robustly powerful than OMiRKAT. Again, I emphasize that enKern enables visualization in a reduced dimensional coordinate space; hence, it is better interpreted. To summarize, enKern enables a unified statistical inference across multiple kernels with the simultaneous achievement of improved power and visual representation, while other omnibus testing methods have focused only on power improvements.

## 2 Methods

### 2.1 Notations and Models

The human microbiome is the entire ecosystem of all microbes that live in and on the human body. In human microbiome studies, researchers seek to find if microbial features (e.g., operational taxonomic units, amplicon sequence variants, species, or genera) are associated with human health or disease adjusting for covariates (e.g., age, sex).

Suppose that there are *n* study subjects (*i* = 1, …, *n*), *p* microbial features (*j* = 1, …, *p*), and *q* covariates (*k* = 1, …, *q*) with the high-dimensionality of *p* ≫ *n*. Then, let *y* denote the *n* × 1 vector for a human health or disease response, *X* denote the *n* × *p* matrix for microbial features, and *Z* denote the *n* × *q* matrix for covariates. Then, to relate microbial features with a continuous response, we can consider a linear regression model as in Eq. (1),

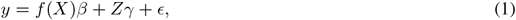

where *f* (·) is a basis function for linear or nonlinear feature mapping; *β* is the *p* × 1 vector of the coefficients on the effects of microbial features: *β* = (*β*_1_, …, *β*_*p*_)^*T*^; *γ* is the *q* × 1 vector of the coefficients on the effects of covariates: *γ* = (*γ*_1_, …, *γ*_*q*_)^*T*^; and *ϵ* is the *n* × 1 vector of independently and identically distributed errors with mean zero and variance *σ*^2^: *ϵ* = (*ϵ*_1_, …, *ϵ*_*n*_)^*T*^. Similarly, for a binary response, we can consider a logistic regression model as in Eq. (2),

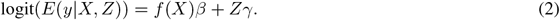

Especially, we are interested in (1) testing the null hypothesis of no effect from all microbial features as in Eq. (3) [1, 2, 3, 32]; and (2) visualizing *p*-dimensional microbial features in a two- or three-dimensional coordinate space.

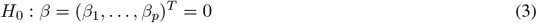

### 2.2 Linear Kernel Method

First of all, for significance testing, the traditional *F*-test or likelihood ratio test cannot be employed because of the high-dimensionality of *p* ≫ *n* and its resulting issues of low rank matrix and low degrees of freedom. Therefore, as in SKAT [1, 2] and MiRKAT [3], we can employ a variance component score test [33, 34]. I describe it using the simplest linear kernel first to survey the linear patterns of the relationship, where the basis function *f* (·) in Eq. (1) and Eq. (2) is the identity (or linear) function, to ease later descriptions on the ecological kernel method (see *Ecological Kernel Method*) and the ensemble learning method, enKern (see *Ensemble Learning Method: enKern*).

We can formulate a test statistic using the linear kernel as

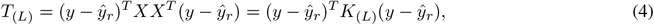

where *ŷ*_*r*_ is an *n* × 1 vector of fitted response values obtained from a null model: *y* = *Zγ*_*r*_ + *ϵ*_*r*_ for Eq (1) and logit(*E*(*y*|*Z*)) = *Zγ*_*r*_ for Eq. (2). Here, the linear kernel *K*_(*L*)_ is an *n* × *n* positive semi-definite matrix that is constructed through the linear kernel function 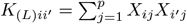 for *i* = 1, …, *n* and *i*^′^ = 1, …, *n*. The linear kernel *K*_(*L*)_ (= *XX*^*T*^) is also the *n* × *n* inner-product (gram) matrix of *X*, which reflects pairwise (i.e., subject-by-subject) similarities in microbial features. We can also see that *y* − *ŷ*_*r*_ is a vector of residuals obtained from the null model, which reflects remaining response values after regressing out the covariate effects on *y*. Interestingly, *T*_(*L*)_ estimates a quantity on the sum of squared microbial effects in a proven sense that SKAT using the linear kernel [1, 2] is equivalent to the sum of squared score tests [35] in the absence of covariates [36, 20]. Then, to obtain null test statistic values that contain only the covariate effects while maintaining the correlation structure across microbial features, we can permute *X* by rows (or, equivalently, the residuals *y* − *ŷ*_*r*_). Then, the *P*-value is calculated as the proportion of null test statistic values,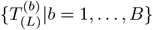, that are greater than or equal to the observed test statistic value, 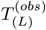 as in Eq. (5).

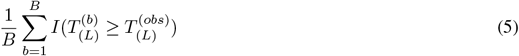

where *I*(·) is an indicator function and *B* is the number of possible permutations.

Note that, in this permutation method, the null model needs to be fitted only once. In contrast, the popular Freedman-Lane permutation method [37] requires to fit the null (or full) model *B* times; that is, the null (or full) model needs to be fitted every time when the residual vector (*y* − *ŷ*_*r*_) is permuted [38, 3]. For the linear regression model, the Freedman-Lane permutation method [37] is widely used since the closed-form least-squares solution is available. However, other regression models, such as the logistic regression model, can require an iterative computational algorithm for maximum likelihood estimation (MLE). Then, the Freedman-Lane permutation method [37] can encounter an error of no convergence in MLE. For this reason, a parametric bootstrap method can be employed as an alternative as in [3]. However, the permutation method described here can flexibly apply to various regression models.

For visual representations, we can construct the singular value decomposition of *X* as in Eq. (6).

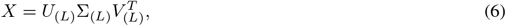

where *U*_(*L*)_ is an *n* × *p* orthogonal matrix with the left eigenvectors as its columns; *V*_(*L*)_ is an *p* × *p* orthogonal matrix with the right eigenvectors as its columns; and Σ_(*L*)_ is an *p* × *p* diagonal matrix with the ordered eigenvalues (i.e., *λ*_(*L*)1_≥ · · · ≥ *λ*_(*L*)*p*_) as its diagonal elements. Then, the first two or three principal components (i.e., the first two or three columns of *U*_(*L*)_Σ_(*L*)_) represent the linear approximations of microbial features with two or three ranks, which in turn enables visual representations in a two- or three-dimensional coordinate space. Here, the *n* non-negative eigenvalues can also describe the proportion of total variance explained by each principal component. Alternatively, we can construct the singular value decomposition of the kernel *K*_(*L*)_ as in Eq. (7)

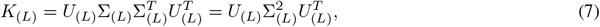

where *U*_(*L*)_ is an *n* × *n* orthogonal matrix with the left eigenvectors as its columns; and Σ_(*L*)_ is an *n* × *n* diagonal matrix with the ordered non-negative eigenvalues (i.e., *λ*_(*L*)1_ ≥ · · · ≥ *λ*_(*L*)*n*_) as its diagonal elements. Then, again, we can use the first two or three principal components (i.e., the first two or three columns of *U*_(*L*)_Σ_(*L*)_) for visual representations, while describing their proportions of total variance explained using their eigenvalues as 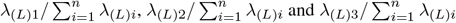 [15, 16, 17, 39].

### 2.3 Ecological Kernel Method

In human microbiome studies, ecological kernels are commonly used to better reflect pairwise (i.e., subject-by-subject) similarities in microbiome composition, which is ultimately to improve statistical power in KAT [3, 8, 9, 10, 11].

An ecological kernel is an *n* × *n* positive semi-definite matrix that is constructed by transforming an ecological distance measure, also known as *β*-diversity index, to a similarity measure through the formula, Eq. (8).

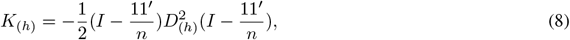

where *h* is an index for a chosen ecological distance measure: 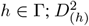 is an *n* × *n* element-wise squared ecological distance matrix; *I* is an *n* × *n* identity matrix; and 1 is an *n* × 1 vector of ones. By its formula Eq. (8), we can infer that a chosen ecological distance measure determines its kernel’s characteristics. The most commonly used ecological distance measures are Jaccard distance [40], Bray-Curtis dissimilarity [41], unweighted UniFrac distance [42], generalized UniFrac distance [43] and weighted UniFrac distance [44].

We can categorize ecological kernels, that is, Jaccard, Bray-Curtis, unweighted UniFrac, generalized UniFrac and weighted UniFrac kernels, into (1) non-phylogenetic kernels (Jaccard and Bray-Curtis kernels) vs. phylogenetic kernels (unweighted UniFrac, generalized UniFrac and weighted UniFrac kernels); and (2) presence-absence kernels (Jaccard and unweighted UniFrac kernels) vs. abundance kernels (Bray-Curtis, generalized UniFrac and weighted UniFrac kernels). We can also describe their characteristics as follows.

First, phylogenetic kernels incorporate phylogenetic tree information, while non-phylogenetic kernels do not. Hence, phylogenetic kernels can be powerful in KAT when the underlying causal variants are phylogenetically close, but it is vice versa when they are not phylogenetically close. Second, presence-absence kernels consider only the dichotomous information (0: absence, 1: presence), while abundance kernels consider the full spectrum of microbial abundance. Hence, presence-absence kernels can be powerful in KAT when the underlying causal variants are rare variants in a nonlinear discrete relationship, but it is vice versa when they are common variants [3]. Finally, the generalized UniFrac kernel [43] has a tuning parameter (0 ≤*α* ≤1) to modulate the extent of abundance information to be incorporated. That is, if *α* is small, it is similar to the unweighted UniFrac kernel, while if *α* is large, it is similar to the weighted UniFrac kernel.

Now, we have various ecological kernels, *K*_(*h*)_’s, that reflect pairwise (i.e., subject-by-subject) similarities in microbial features with different views. Recall from Eq. (4) that we can think of the linear kernel *K*_(*L*)_ (= *XX*^*T*^) as the *n* × *n* inner-product (gram) matrix of the *n* × *p* original microbial feature matrix *X*. However, there is no underlying *p*-dimensional microbial feature matrix: *f*_(*h*)_(*X*), where *f*_(*h*)_(·) is possibly a nonlinear basis function, that can be directly observed from *K*_(*h*)_ (= *f*_(*h*)_(*X*)*f*_(*h*)_(*X*)^*T*^). The reason is because we constructed the ecological kernel, *K*_(*h*)_, through the conversion from distance to similarity in Eq. (8), not explicitly specifying the basis function, *f*_(*h*)_(·). However, the Mercer’s theorem [45] ensures that even when *f*_(*h*)_(*X*) is unknown, we can obtain its orthogonal lower-dimensional representations, *X*_(*h*)_, through the singular value decomposition of *K*_(*h*)_ as in Eq. (9).

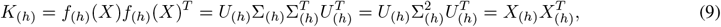

where *U*_(*h*)_ is an *n* × *n* orthogonal matrix with the left eigenvectors as its columns; Σ_(*h*)_ is an *n* × *n* diagonal matrix with the ordered non-negative eigenvalues (i.e., *λ*_(*h*)1_ ≥ · · · ≥ *λ*_(*h*)*n*_) as its diagonal elements; and *X*_(*h*)_ = *U*_(*h*)_Σ_(*h*)_ for *h* ∈ Γ. Here, we can see that 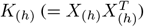 is the *n* × *n* inner-product (gram) matrix of the *n* × *n* principal component matrix *X*_(*h*)_ that is referred here as the surrogate microbial feature matrix. Then, the principal components (i.e., the columns of *X*_(*h*)_) can be referred as surrogate microbial features. To be more detailed, the surrogate microbial features belong to a lower-dimensional space of the reproducing kernel Hilbert space for a kernel *K*_(*h*)_ [45]. Then, we can formulate a test statistic using an ecological kernel as in Eq. (10).

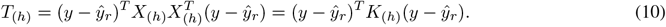

For KAT, as in the linear kernel method, we can permute *X*_(*h*)_ by rows (or, equivalently, the residuals *y* − *ŷ*_*r*_) to obtain null test statistics values. Then, the *P*-value is calculated as the proportion of null test statistic values, 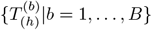, that are greater than or equal to the observed test statistic value, 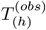 as in Eq. (11).

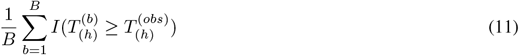

where *I*(·) is an indicator function and *B* is the number of possible permutations.

For PCA, as we did in Eq. (7) for the linear kernel method, we can employ the singular value decomposition of the kernel *K*_(*h*)_ in Eq. (9). Then, for visual representations, we can pick the first two or three principal components (i.e., the first two or three columns of *U*_(*h*)_Σ_(*h*)_), while describing their proportions of total variance explained using their eigenvalues 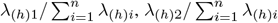 and 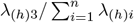 [15, 16, 17, 39].

### 2.4 Ensemble Learning Method: enKern

We can so far conduct KAT and PCA jointly on each kernel; yet, the choice of kernel can dramatically alter the outcomes in both KAT and PCA. In this section, I describe methodological details on enKern to conduct KAT and PCA jointly on multiple kernels. All the computational procedures are summarized in Algorithm: enKern.

I first formulate a weighted test statistic as in Eq. (12).

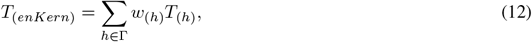

where *T*_(*h*)_ is the test statistic for each kernel *h* in Eq. (10); Γ is a set of all candidate kernels; and *w*_(*h*)_ is the weight for each test statistic. Here, the weight for each test statistic is the reciprocal of its standard deviation under *H*_0_ as in Eq. (13)

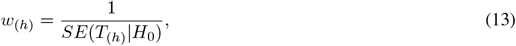

where *SE*(*T*_(*h*)_|*H*_0_) is estimated by the standard deviation of null (permuted) test statistic values: *SD* 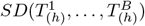 adjusted to have Σ _*h*∈ Γ_ *w*_(*h*)_ = 1. Note that to account for the correlations across individual kernel association tests, the rows of *X*_(*h*)_’s should be permuted simultaneously to have the same sequence of rearrangements across all candidate kernels in Γ. Alternatively, we can apply the same permuted residuals *y* − *ŷ*_*r*_ to all candidate kernels in Γ simultaneously. By Eq. (12) and Eq. (13), we can infer that test statistics that have a large effect (i.e., a large *T*_(*h*)_ value) and a small standard error (i.e., a small *SE*(*T*_(*h*)_ | *H*_0_) value) will be attached more to the weighted test statistic of enKern, *T*_(*enKern*)_. To explain its rationale, it is first obvious that a large effect (i.e., a large *T*_(*h*)_ value) gives more credence against *H*_0_ (i.e., toward *H*_1_) [a small effect (i.e., a small *T*_(*h*)_ value) gives more credence against *H*_1_ (i.e., toward *H*_0_)]. However, if its standard error is large under *H*_0_ (i.e., a large *SE*(*T*_(*h*)_ | *H*_0_) value), even a large effect (i.e., a large *T*_(*h*)_ value) will be likely to occur under *H*_0_ [if its standard error is small under *H*_0_ (i.e., a small *SE*(*T*_(*h*)_ | *H*_0_) value), even a small effect (i.e., a small *T*_(*h*)_ value) will be less likely to occur under *H*_0_]. Therefore, a large standard error (i.e., a large *SE*(*T*_(*h*)_ | *H*_0_) value) gives more credence toward *H*_0_ (i.e., against *H*_1_) [a small standard error (i.e., a small *SE*(*T*_(*h*)_ | *H*_0_) value) gives more credence toward *H*_1_ (i.e., against *H*_0_)].

It is fundamentally different from the popular minimum *P*-value approach. Its minimum *P*-value test statistic reflects only the strongest association signal; yet, the weighted test statistic of enKern leverages complementary contributions from multiple input kernels attaching strong association signals but suppressing weak association signals. Thus, enKern can better adapt to various sparsity levels, while the minimum *P*-value approach is powerful for a high sparsity level, where only few tests among multiple association tests make strong association signals.

Another critical advantage of enKern over the minimum *P*-value approach is in its computational efficiency. That is, it takes only a second to calculate its *P*-value given all the observed 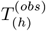 and null 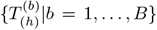 test statistic values for all individual candidate kernels for *h* ∈ Γ. In other words, once all individual kernel association tests are done, there is almost no additional computational burden to calculate the *P*-value for enKern; hence, overall computational burden increases only linearly as the number of candidate kernels increases. Importantly, this promises potential extensions to more kernels in a computationally efficient manner.

To be more precise, to calculate the *P*-value for enKern, we first need to calculate the weight for each test statistic as 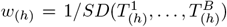 for *h* ∈ Γ, which is a simple non-iterative arithmetic operation given 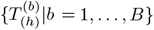 for *h* ∈ Γ. Then, we need to calculate the *P*-value for enKern as the proportion of null test statistic values, 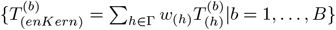, that are greater than or equal to the observed test statistic value, 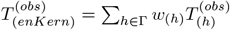 as in Eq. (14), which is again a simple non-iterative arithmetic operation given *w*_(*h*)_, 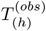 and 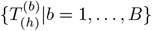 for *h* ∈ Γ.

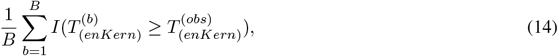

where *I*(·) is an indicator function and *B* is the number of possible permutations. On the contrary, the minimum *P*-value approach requires double (e.g., *B* × *B*) iterations because the minimum *P*-value itself is a test statistic for the minimum *P*-value approach. That is, to calculate a *P*-value, we need to calculate *B* null test statistic values; yet again to calculate a *P*-value based on the minimum *P*-value test statistic, we need to calculate *B* null minimum *P*-value test statistic values.

For PCA, I define a weighted kernel as in Eq. (15).

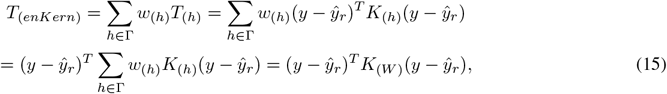

where *K*_(*W*)_ (= Σ _*h*∈ Γ_ *w*_(*h*)_ *K*_(*h*)_) is the weighted kernel. Here, we can see that it is mathematically equivalent to apply the weights (i.e., *w*_(*h*)_’s) to individual test statistics (i.e., *T*_(*h*)_’s) or individual kernels (i.e., *K*_(*h*)_’s) to formulate the weighted test statistic (i.e., *T*_(*enKern*)_). Then, we can also conduct significance testing for enKern through the singular value decomposition of the kernel *K*_(*W*)_ as in Eq. (16) based on the test statistic using the weighted kernel as in Eq. (17) because *T*_(*enKern*)_ in Eq. (15) is equivalent to *T*_(*W*)_ in Eq. (17).

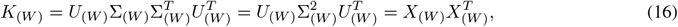

where *U*_(*W*)_ is an *n* × *n* orthogonal matrix with the left eigenvectors as its columns; Σ_(*W*)_ is an *n* × *n* diagonal matrix with the ordered non-negative eigenvalues (i.e., *λ*_(*W*)1_ ≥ · · · ≥ *λ*_(*W*)*n*_) as its diagonal elements; and *X*_(*W*)_ = *U* (*W*) Σ (*W*).

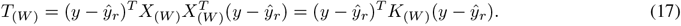

Note also that the weighted kernel *K*_(*W*)_ is also a valid positive semi-definite kernel because it is a linear combination of positive semi-definite matrices: Σ _*h*∈ Γ_ *w*_(*h*)_ *K*_(*h*)_. Importantly, the singular value decomposition of the weighted kernel *K*_(*W*)_ in Eq. (16) facilitates dimension reduction and visualization. That is, as we did in Eq. (7) for the linear kernel method and in Eq. (9) for the ecological kernel, we can employ the singular value decomposition of the weighted kernel *K*_(*W*)_ in Eq. (16). Then, we can pick the first two or three principal components (i.e., the first two or three columns of *U*_(*W*)_Σ_(*W*)_), while describing their proportions of total variance explained using their eigenvalues as ^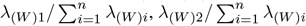 and 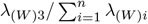^ In this way, we can conduct dimension reduction and visualization to be perfectly matched with their original significance testing scheme as in Eq. (15) and Eq. (17).

#### Algorithm

enKern

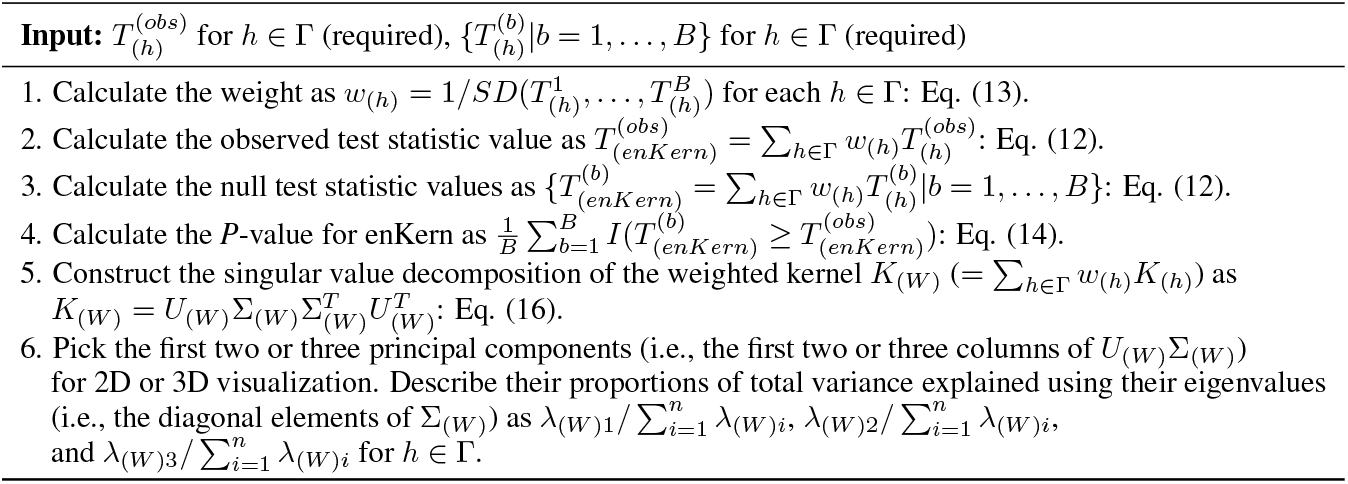

## 3 Results

To demonstrate the use of enKern, I surveyed (1) the disparity in upper-respiratory-tract microbiome by cigarette smoking as in [30, 3]; and (2) the disparity in oral (subgingival) microbiome by gingival inflammation and continuous cytokine (IL-8) levels, respectively, as in [31]. I employed the Jaccard, Bray-Curtis, unweighted UniFrac, generalized UniFrac (0.25), generalized UniFrac (0.5), generalized UniFrac (0.75) and weighted UniFrac kernels because of their unique and well-distinguished characteristics as well as their popularity in human microbiome *β*-diversity analysis.

### 3.1 The disparity in upper-respiratory-tract microbiome by cigarette smoking

The upper-respiratory-tract microbiome data contain 857 microbial features for 60 study subjects, but I removed microbial features that have the mean proportion < .00005 and three study subjects that had have antibiotic treatment within the last three months; as such, 402 microbial features (*p* = 402) for 57 study subjects (*n* = 57) were finally retained. I added age and sex as covariates in the analysis. I surveyed the smoking status as binary (31 nonsmokers and 26 smokers) and continuous traits (the number of packs of cigarettes per year), respectively.

First, I found a significant disparity in upper-respiratory-tract microbiome composition by smoking status (31 nonsmokers and 26 smokers) using enKern (*P*-value: 0.00135) and OMiRKAT [3] (*P*-value: 0.00220) at the significance level of 0.05, yet we can see that the *P*-value for enKern is smaller than the *P*-value for OMiRKAT [3] (Table 2A and Table 2B). Accordingly, we can also visually see a clear distinction in upper-respiratory-tract microbiome composition that nonsmokers (colored in blue) and smokers (colored in red), respectively, tend to be located closely to each other in the PCA plots (Figure 1A).

**Table 1:** The *P*-values from the analyses using individual kernel and omnibus testing or ensemble learning methods on (**A**) the disparity in upper-respiratory-tract microbiome composition by smoking status; (**B**) the disparity in upper-respiratory-tract microbiome composition by the numbers of packs of cigarettes per year; (**C**) the disparity in oral microbiome composition by gingival inflammation status; (**D**) the disparity in oral microbiome composition by cytokine (IL-8) levels. \**K*_*J*_, *K*_*BC*_, *K*_*U*_, *K*_0.25_, *K*_0.5_, *K*_0.75_ and *K*_*W*_ represent the use of Jaccard, Bray-Curtis, unweighted UniFrac, generalized UniFrac (0.25), generalized UniFrac (0.5), generalized UniFrac (0.75) and weighted UniFrac kernels, respectively.

| Kernel | A | B | C | D |
| --- | --- | --- | --- | --- |
| $K_J$ | 0.00305 | 0.03110 | 0.12554 | 0.73532 |
| $K_{BC}$ | 0.00071 | 0.03279 | 0.05938 | 0.94490 |
| $K_U$ | 0.00195 | 0.06046 | 0.86093 | 0.65833 |
| $K_{0.25}$ | 0.00196 | 0.05166 | 0.01335 | 0.96340 |
| $K_{0.5}$ | 0.00207 | 0.04818 | 0.01333 | 0.97500 |
| $K_{0.75}$ | 0.00277 | 0.05692 | 0.01733 | 0.97970 |
| $K_W$ | 0.00420 | 0.07733 | 0.02655 | 0.98110 |
| OMiRKAT | 0.00220 | 0.06923 | 0.04123 | 0.94820 |
| enKern | 0.00135 | 0.04113 | 0.02704 | 0.97100 |

**Table 2:** The empirical type 1 error rates for individual kernel and omnibus testing or ensemble learning methods (Unit: %). \**K*_*J*_, *K*_*BC*_, *K*_*U*_, *K*_0.25_, *K*_0.5_, *K*_0.75_ and *K*_*W*_ represent the use of Jaccard, Bray-Curtis, unweighted UniFrac, generalized UniFrac (0.25), generalized UniFrac (0.5), generalized UniFrac (0.75) and weighted UniFrac kernels, respectively.

| Continuous |  |  | Binary |  |  |
| --- | --- | --- | --- | --- | --- |
| Kernel | $n = 100$ | $n = 200$ | Kernel | $n = 100$ | $n = 200$ |
| $K_J$ | 5.02 | 4.84 | $K_J$ | 4.88 | 4.98 |
| $K_{BC}$ | 4.87 | 5.17 | $K_{BC}$ | 4.75 | 4.71 |
| $K_U$ | 4.92 | 4.96 | $K_U$ | 4.92 | 4.79 |
| $K_{0.25}$ | 5.12 | 4.92 | $K_{0.25}$ | 4.75 | 5.09 |
| $K_{0.5}$ | 5.00 | 4.92 | $K_{0.5}$ | 5.04 | 4.75 |
| $K_{0.75}$ | 5.00 | 4.96 | $K_{0.75}$ | 5.04 | 4.92 |
| $K_W$ | 5.04 | 4.71 | $K_W$ | 5.00 | 5.04 |
| OMiRKAT | 5.01 | 4.98 | OMiRKAT | 5.00 | 4.97 |
| enKern | 4.96 | 5.02 | enKern | 4.98 | 4.98 |

**Figure 1.**
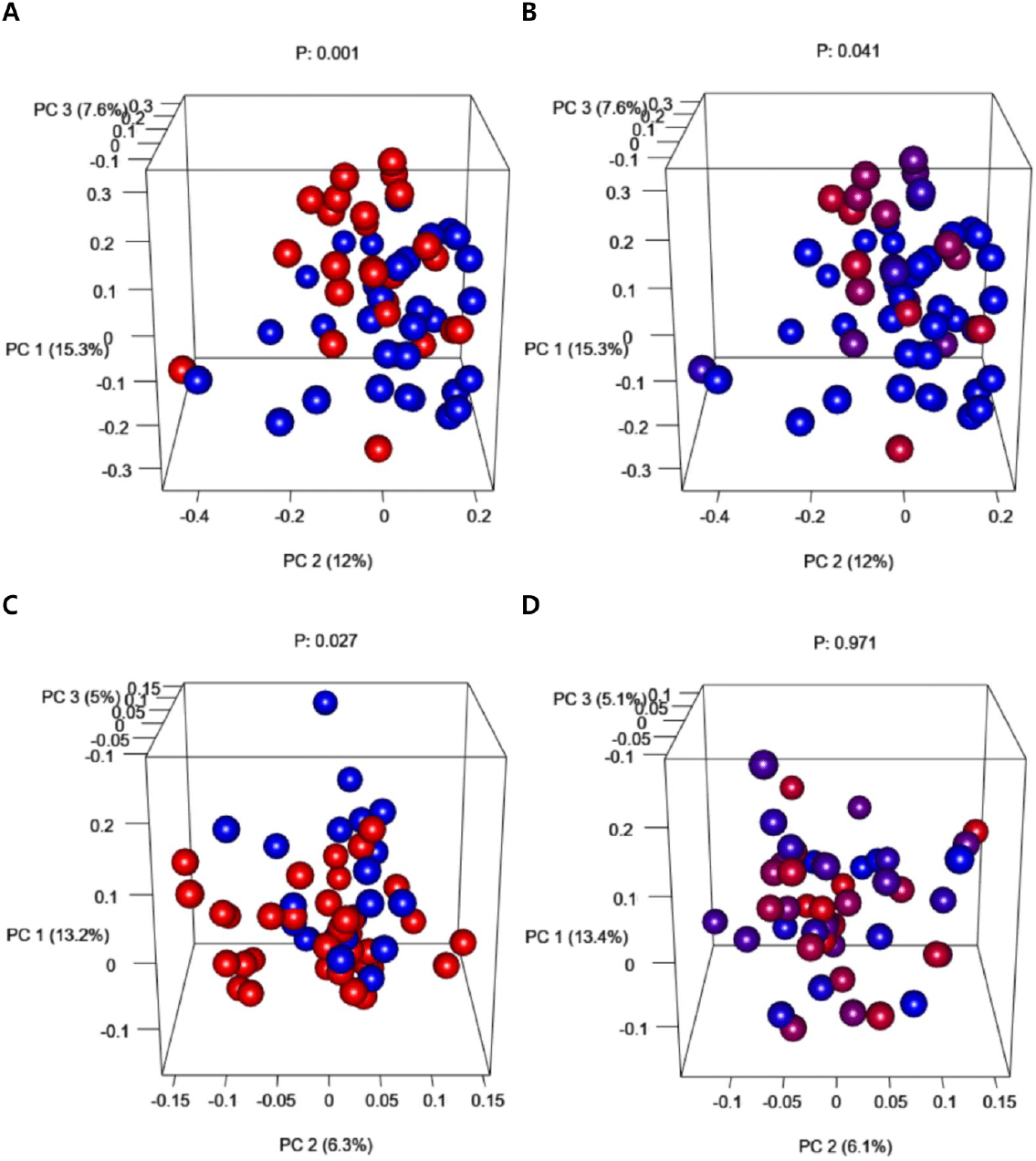
The three dimensional PCA plots to demonstrate (**A**) the disparity in upper-respiratory-tract microbiome composition by smoking status: 31 nonsmokers (colored in blue) and 26 smokers (colored in red); (**B**) the disparity in upper-respiratory-tract microbiome composition by the numbers of packs of cigarettes per year: gradient colors ranging from the minimum 0 pack of cigarettes per year to be colored in blue to the maximum 42 packs of cigarettes per year to be colored in red; (**C**) the disparity in oral microbiome composition by gingival inflammation status: 22 healthy individuals (colored in blue) and 35 individuals with gingival inflammation (colored in red); (**D**) the disparity in oral microbiome composition by cytokine (IL-8) levels: gradient colors ranging from the minimum 89.19 IL-8 level to be colored in blue to the maximum 2155.48 IL-8 level to be colored in red. *P represents the *P*-value using enKern. *The three axes represent the first three principal components with their percentages of total variance explained using enKern.

Second, I found a significant disparity in upper-respiratory-tract microbiome composition by the numbers of packs of cigarettes per year using enKern (*P*-value: 0.04113) at the significance level of 0.05, yet it was not significant for OMiRKAT [3] (*P*-value: 0.06923) (Table 2A and Table 2B). Accordingly, we can also visually see a clear distinction in upper-respiratory-tract microbiome composition that the colors close to blue for the minimum 0 pack of cigarettes per year and the colors close to red for the maximum 42 packs of cigarettes per year, respectively, tend to be located closely to each other in the PCA plots (Figure 1B).

### 3.2 The disparity in oral microbiome by gingival inflammation status and cytokine levels

The oral microbiome data contain 2,261 microbial features for 57 study subjects that are male e-cigarette users measured at the initial recruitment [31], but I removed microbial features that have the mean proportion < .00005; as such, 1,486 microbial features (*p* = 1, 486) for 57 study subjects (*n* = 57). I added age and the frequency of brushing teeth as covariates in the analysis. First, I found a significant disparity in oral microbiome composition by gingival inflammation status (22 healthy individuals and 35 individuals with gingival inflammation) using enKern (*P*-value: 0.02704) and OMiRKAT [3] (*P*-value: 0.04123) at the significance level of 0.05, yet we can see that the *P*-value for enKern is smaller than the *P*-value for OMiRKAT [3] (Table 2C and Table 2D). Accordingly, we can also visually see a clear distinction in oral microbiome composition that healthy individuals (colored in blue) and individuals with gingival inflammation (colored in red), respectively, tend to be located closely to each other in the PCA plots (Figure 1C).

Second, I could not find a significant disparity in oral microbiome composition by cytokine (IL-8) levels using enKern (*P*-value: 0.97100) and OMiRKAT [3] (*P*-value: 0.94820) at the significance level of 0.05 (Table 2C and Table 2D). Accordingly, we cannot also see any visual distinction in oral microbiome composition by cytokine (IL-8) levels that the colors close to blue for the minimum 89.19 IL-8 level and the colors close to red for the maximum 2155.48 IL-8 level are all randomly spread in the PCA plots (Figure 1D).

### 3.3 Simulation Designs

As in prior studies [3, 23], I conducted simulation studies to evaluate the performance of enKern compared with OMiRKAT [3] in significance testing. For this, I began with the Charlson et al.’s upper-respiratory-tract microbiome data [30] that I used for my first real data application. Then, to reflect real microbiome composition, I estimated their mean proportions and over-dispersion based on the Dirichlet-multinomial distribution [46]. Then, I generated random counts for 100 (*n* = 100) and 200 (*n* = 200) subjects, respectively, with the library size of 10,000 from the Dirichlet-multinomial distribution [46] on the estimated mean proportions and over-dispersion. Then, I generated (1) continuous responses based on the linear regression model,

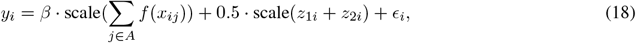

and (2) binary responses based on the logistic regression model,

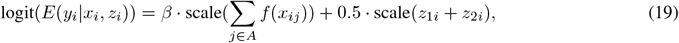

where *f* (·) is the standardization function to have mean 0 and variance 1 for each variant; *A* is a set of microbial features that can be related to the response *y*_*i*_ through the regression coefficient *β*; *z*_1*i*_ and *z*_2*i*_ are covariates generated to be independent with microbial features as *z*_1*i*_ ~ *Bern*(0.5) and to be correlated with 10% of randomly selected microbial features (*C*) as *z*_2*i*_ ~ 0.5 × scale(Σ _*j*∈*C*_ *x*_*ij*_) + *N* (0, 1), respectively; *ϵ*_*i*_ is a random error generated as *ϵ*_*i*_ ~ *N* (0, 1); and scale is the standardization function to have mean 0 and variance 1.

To evaluate type I error rate, I set *β* = 0. To evaluate power, I set *β* = 2 for Eq. (18) and *β* = 4 for Eq. (19). In the meanwhile, I surveyed four different association scenarios while differing the microbial features to be included in *A* as: (1) 10% rarest microbial features; (2) 10% randomly selected microbial features; (3) 10% most common microbial features; and (4) microbial features in one lineage among ten phylogenetic lineages, respectively. The first three scenarios mimic abundance-based association patterns, where rare, random selected or common microbial features are related to the response. The fourth scenario mimics phylogeny-based association patterns, where phylogenetically close microbial features are related to the response. To be more specific, for the fourth scenario, I partitioned microbial features into ten lineages using the partitioning-around-medoids algorithm [47] based on microbial features’ cophenetic distances [48]. Then, I set the microbial features in the rarest (occupied 5.2% of total abundance), medium (occupied 9.9% of total abundance) or the most common (occupied 19.8% of total abundance) lineage to be related to the response.

### 3.4 Simulation Results

#### 3.4.1 Type 1 error

I organized the empirical type 1 error rates in Table 2. We can see that the type 1 error rates are well-controlled at the significance level of 0.05 for all individual kernel and omnibus testing or ensemble learning methods for both sample sizes of (*n* = 100) and (*n* = 200), and also for both continuous and binary responses (Table 2).

#### 3.4.2 Power

I organized the empirical power values in Figure 2. I also organized them in Table 3 for the continuous responses and in Table 4 for the binary responses using numbers. We can first see that the power values are larger when the sample size is larger: Figure 2A < Figure 2B; Figure 2C < Figure 2D, as expected. We can also see that the power values are larger for the continuous responses than the binary responses: Figure 2C < Figure 2A; Figure 2D < Figure 2B, as expected. However, note that the relative (not absolute) performances across individual kernel and omnibus testing or ensemble learning methods are crucial. We can see that the relative performances are maintained similarly for each of the abundance-based association scenarios for the rare (A1), randomly selected (A2) and common (A3) microbial features, and the phylogeny-based association scenarios for the rare (P1), medium (P2) and common (P3) lineages (Figure 2). I dissected it further in details in the following paragraph.

**Table 3:** The empirical power values for individual kernel and omnibus testing or ensemble learning methods for the continuous responses (Unit: %). \***A1, A2** and **A3** represent the abundance-based association scenarios, where the rare (**A1**), random selected (**A2**) or common (**A3**) microbial features are related to the response. \***P1, P2** and **P3** represent the phylogeny-based association scenarios, where the microbial features in the rare (**P1**), medium (**P2**) or common (**P3**) phylogenetic lineage are related to the response. \**K*_*J*_, *K*_*BC*_, *K*_*U*_, *K*_0.25_, *K*_0.5_, *K*_0.75_ and *K*_*W*_ represent the use of Jaccard, Bray-Curtis, unweighted UniFrac, generalized UniFrac (0.25), generalized UniFrac (0.5), generalized UniFrac (0.75) and weighted UniFrac kernels, respectively.

| Continuous response ($n = 100$) | | | | | | |
| --- | --- | --- | --- | --- | --- | --- |
| Kernel | A1 | A2 | A3 | P1 | P2 | P3 |
| $K_J$ | 13.57 | 10.21 | 8.55 | 8.90 | 8.20 | 8.80 |
| $K_{BC}$ | 9.35 | 22.75 | 57.90 | 11.65 | 19.20 | 15.50 |
| $K_U$ | 11.99 | 10.53 | 7.80 | 6.95 | 10.80 | 8.00 |
| $K_{0.25}$ | 10.46 | 17.94 | 37.40 | 52.17 | 45.01 | 54.28 |
| $K_{0.5}$ | 10.46 | 17.75 | 40.90 | 65.25 | 53.56 | 65.97 |
| $K_{0.75}$ | 10.03 | 17.36 | 38.35 | 68.9 | 55.65 | 66.80 |
| $K_W$ | 9.18 | 16.06 | 32.35 | 67.96 | 55.54 | 64.75 |
| OMiRKAT | 10.80 | 15.48 | 40.13 | 48.58 | 39.94 | 48.62 |
| enKern | 11.48 | 20.41 | 41.30 | 52.80 | 46.25 | 53.70 |
| Continuous response ($n = 200$) | | | | | | |
| Kernel | A1 | A2 | A3 | P1 | P2 | P3 |
| $K_J$ | 17.38 | 19.21 | 11.76 | 12.51 | 12.82 | 13.12 |
| $K_{BC}$ | 7.69 | 51.84 | 95.36 | 28.57 | 47.64 | 32.04 |
| $K_U$ | 14.10 | 13.82 | 11.38 | 9.88 | 15.98 | 12.29 |
| $K_{0.25}$ | 10.64 | 40.57 | 79.68 | 95.25 | 85.96 | 94.26 |
| $K_{0.5}$ | 10.51 | 41.75 | 83.38 | 97.84 | 92.51 | 97.08 |
| $K_{0.75}$ | 10.51 | 37.53 | 80.59 | 97.55 | 93.25 | 96.57 |
| $K_W$ | 10.38 | 32.73 | 72.60 | 96.72 | 93.36 | 94.93 |
| OMiRKAT | 11.53 | 36.82 | 85.40 | 92.42 | 84.89 | 89.38 |
| enKern | 12.05 | 46.35 | 84.36 | 95.21 | 89.18 | 93.25 |

**Table 4:**
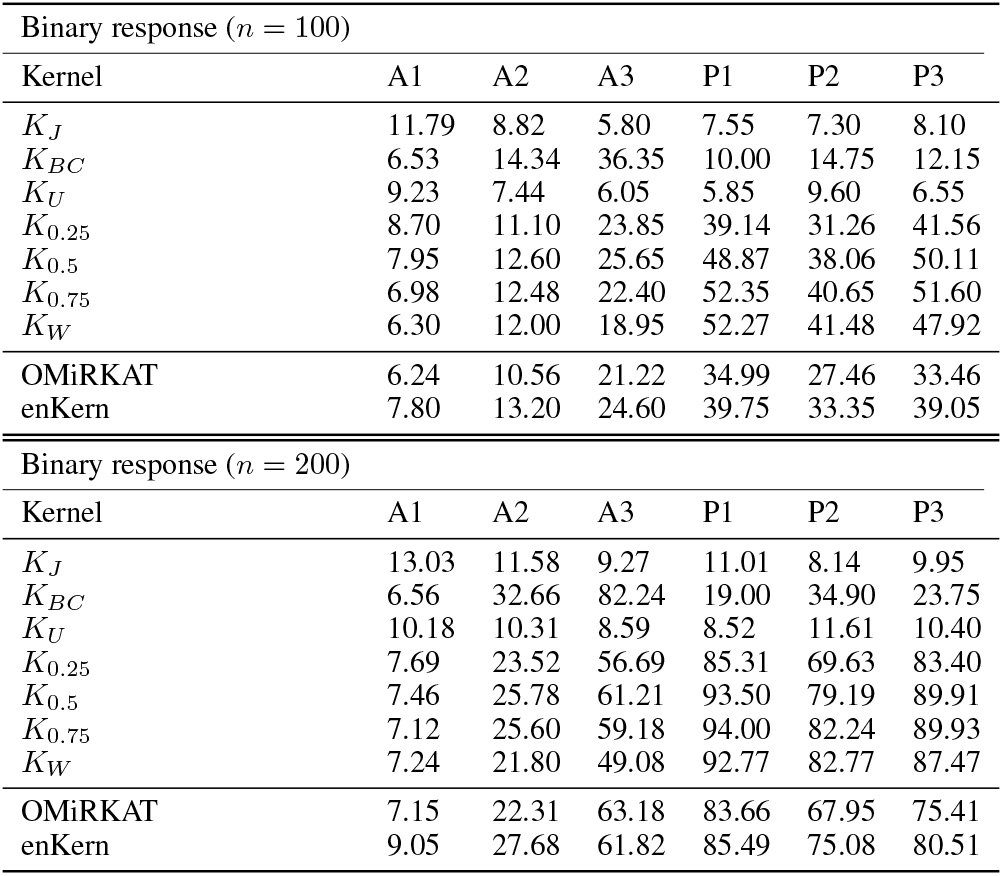
The empirical power values for individual kernel and omnibus testing or ensemble learning methods for the binary responses (Unit: %). \***A1, A2** and **A3** represent the abundance-based association scenarios, where the rare (**A1**), random selected (**A2**) or common (**A3**) microbial features are related to the response. \***P1, P2** and **P3** represent the phylogeny-based association scenarios, where the microbial features in the rare (**P1**), medium (**P2**) or common (**P3**) phylogenetic lineage are related to the response. \**K*_*J*_, *K*_*BC*_, *K*_*U*_, *K*_0.25_, *K*_0.5_, *K*_0.75_ and *K*_*W*_ represent the use of Jaccard, Bray-Curtis, unweighted UniFrac, generalized UniFrac (0.25), generalized UniFrac (0.5), generalized UniFrac (0.75) and weighted UniFrac kernels, respectively.

**Figure 2.**
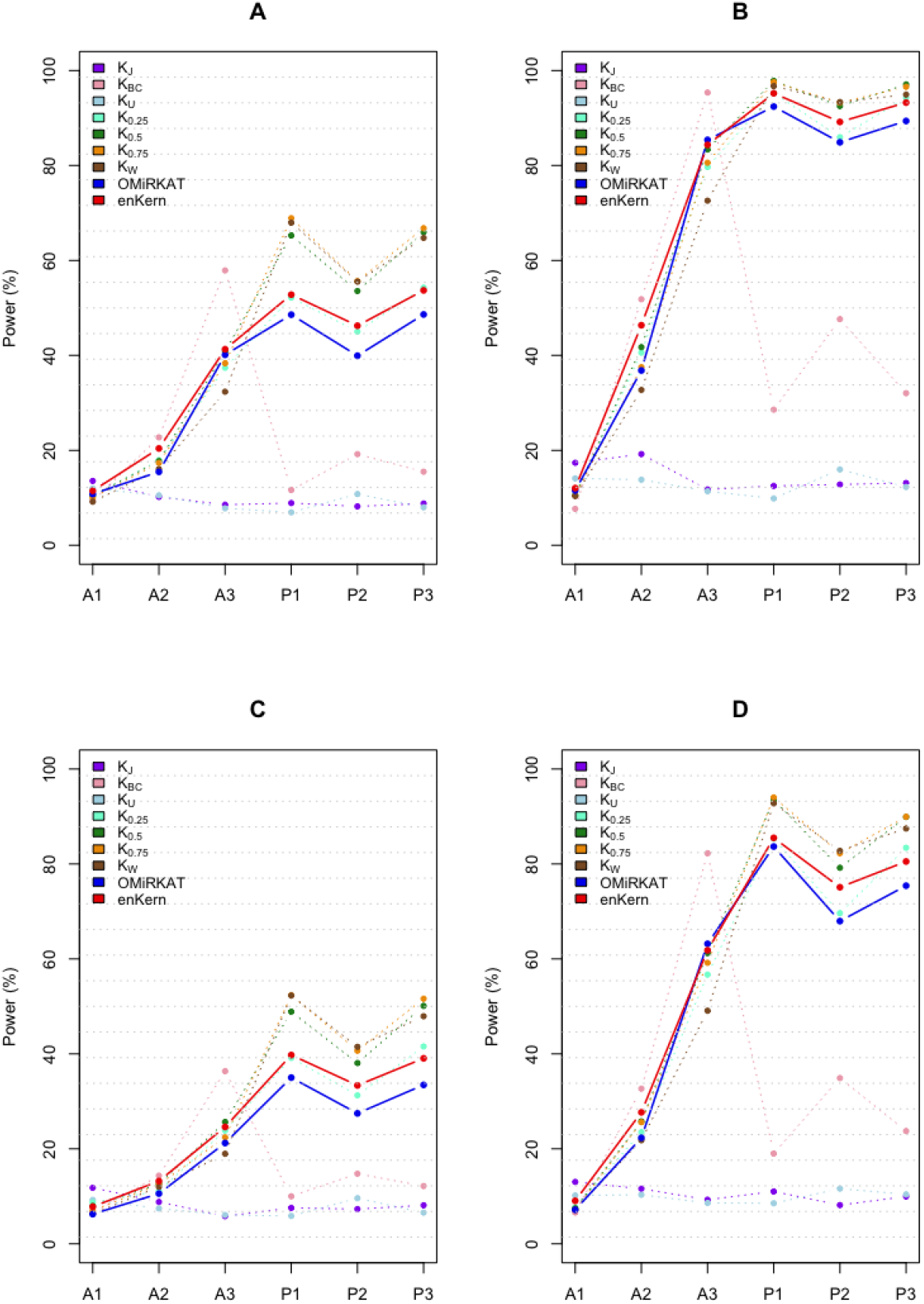
The empirical power values for individual kernel and omnibus testing or ensemble learning methods (Unit: %). \***A** is for continuous responses (*n* = 100); **B** is for continuous responses (*n* = 200); **C** is for binary responses (*n* = 100); **D** is for binary responses (*n* = 200). \***A1, A2** and **A3** represent the abundance-based association scenarios, where the rare (**A1**), random selected (**A2**) or common (**A3**) microbial features are related to the response. \***P1, P2** and **P3** represent the phylogeny-based association scenarios, where the microbial features in the rare (**P1**), medium (**P2**) or common (**P3**) phylogenetic lineage are related to the response. \**K*_*J*_, *K*_*BC*_, *K*_*U*_, *K*_0.25_, *K*_0.5_, *K*_0.75_ and *K*_*W*_ represent the use of Jaccard, Bray-Curtis, unweighted UniFrac, generalized UniFrac (0.25), generalized UniFrac (0.5), generalized UniFrac (0.75) and weighted UniFrac kernels, respectively.

We can see that the power values are larger for the Jaccard kernel (*K*_*J*_) or the unweighted UniFrac kernel (*K*_*U*_) when the rare microbial features are related (A1) (Figure 2, Table 3, Table 4), which is because they consider only the dichotomous (presence-absence) information; yet they lose power for the other scenarios (A2, A3, P1, P2, P3) (Figure 2, Table 3, Table 4). The power values are also larger for the Bray-Curtis kernel *K*_*BC*_ when the random selected (A2) or common (A3) microbial features are related (Figure 2, Table 3, Table 4), which is because the Bray-Curtis kernel considers the full spectrum of microbial abundance; yet it loses power for the other scenarios (A1, P1, P2, P3) (Figure 2, Table 3, Table 4). The power values are also larger for the UniFrac kernels, *K*_0.5_, *K*_0.75_ and *K*_*W*_, when phylogenetically close microbial features are related (P1, P2, P3) (Figure 2, Table 3, Table 4), which is because they incorporate phylogenetic tree information to formulate pairwise (i.e., subject-by-subject) similarities, yet they tend to lose power for abundance-based association scenarios (A1, A2, A3) (Figure 2, Table 3, Table 4).

Importantly, we can see that enKern is moderately more powerful than OMiRKAT [3] across all scenarios except for the abundance-based association scenario when the common microbial features are related to the response (A3) with the same size of *n* = 200 (Figure 2, Table 3, Table 4), indicating that the weight learning scheme of enKern works more robustly than focusing only on the strongest association signal in the minimum *P*-value approach. The reason is because not only the strongest association signal is informative, but also the next strongest association signals can help for (A1, A2, P1, P2, P3) (Figure 2, Table 3, Table 4).

## 4 Discussion

In this paper, I introduced an ensemble learning method, enKern, to conduct KAT and PCA jointly on multiple kernels. enKern is based on a weight learning scheme that leverages complementary contributions from multiple kernels attaching informative association signals but suppressing noisy association signals. It is fundamentally different from the minimum *P*-value approach [19] that focuses only on the strongest association signal. I also showed that applying the weights to individual test statistics or individual kernels is mathematically equivalent to formulate its weighted test statistic, which in turn enables dimension reduction and visualization based on the weighted kernel to be perfectly matched with its original significance testing scheme. Therefore, enKern enables a visualization in a reduced dimensional coordinate space, while other omnibus testing methods have focused only on power improvements. I also demonstrated through simulation studies that enKern achieves a robustly high power for various association patterns. To summarize, enKern enables a unified statistical inference across multiple kernels with the simultaneous achievement of improved power and visual representation.

I demonstrated the use of enKern for human microbiome *β*-diversity analysis, yet its application can be much broader. In human genomics, researchers have focused more on improving statistical power because it is too challenging to discover causal genetic variants. However, it does not necessarily mean that interpretability does not matter in human genomics; hence, enKern can also be a promising analytic tool in human genomics. Of course, KAT and PCA can also apply to other high-dimensional omics studies, such as proteomics, transcriptomics, metabolomics, phenomics, and so forth, or medical image data, such as functional magnetic resonance image or positron emission tomography data. I hope enKern to be widely used for any domains, where KAT and PCA are needed.

I set up candidate kernels in my simulations and real data applications using ecological kernels because of their unique features and popularity in human microbiome studies. However, in reality, there are even more kernels, and an appropriate set of candidate kernels can vary by domains. In tradition, the linear, polynomial, gaussian kernels, and so forth have been considered. For another example, SKAT [1, 2] employed the linear, quadratic and identity-by-state kernels, and their weighted versions. Developing new ecological kernels for human microbiome studies can also advance our understanding of microbial ecology and make the best use of enKern. Note again that enKern promises potential extensions to more kernels in a computationally efficient manner as its overall computational burden increases only linearly as the number of candidate kernels increases. Extensions to various types of response variables and/or study designs as in [4, 5, 6, 7, 8, 9, 10, 11] are also needed. Though I could not satisfy all such demands in this study.

## Acknowledgements

The author is grateful to Drs. Xihong Lin, Michael Wu, Seunggeun Lee, Ni Zhao and Jun Chen for their foundational works on kernel association testing.

## Data Availability Statement

I used two public microbiome datasets (1) to see the disparity in upper-respiratory-tract microbiome by cigarette smoking [30]; and (2) to see the disparity in oral microbiome by gingival inflammation and cytokine levels [31]. The first microbiome dataset is available in the R package, GUniFrac (https://cran.r-project.org/web/packages/GUniFrac), with three R objects: throat.otu.table, throat.tree and throat.meta. The second microbiome dataset is available in the R package, enKern (https://github.com/hk1785/enkern), with three R objects: oral.otu.table, oral.tree and oral.meta.

enKern is freely available in the R package, enKern (https://github.com/hk1785/enkern). All the detailed instructions on the inputs, outputs, arguments and options with example data can also be found in the software webpage.

